# Resveratrol increases tear production and spontaneous ocular pain after corneal abrasion in male, but not female, rats using a preclinical model of photorefractive keratectomy (PRK)

**DOI:** 10.1101/2022.01.25.477730

**Authors:** Deborah M. Hegarty, James R. Carroll, Dennis Nguyen, Victoria S. Halls, Dennis I. Robbins, Theodore J. Price, Gregory Dussor, Sue A. Aicher

**Author notes:** Corresponding author Sue A. Aicher, Ph.D.

## Abstract

Photorefractive keratectomy (PRK) is an alternative to LASIK and can cause intense acute pain that is often not relieved by standard treatments. To assess potential therapeutics for this type of acute pain, appropriate preclinical models are needed. Herein we describe a rodent preclinical model of PRK and a multi-faceted approach to determine the therapeutic potential of resveratrol, a natural phytoestrogen, on pain, tear production, and the corneal epithelium. Studies were conducted in male and female Sprague-Dawley rats. Heptanol was applied to one eye and the superficial corneal epithelium was removed, mimicking the abrasion seen in PRK. Spontaneous pain was assessed with orbital tightening (OT) scores for 7 days. Corneal abrasion increased OT scores in both male and female rats with peak responses at 24 - 48 hours. Topical application of resveratrol had a sex-specific effect on OT scores and tear production. Resveratrol increased OT scores in abraded males, but not females, at 72 hours and 1 week after abrasion. Resveratrol dose-dependently increased tear production in abraded males, but had no effect in abraded females. While there was no correlation between OT score at 1 week and tear production, CGRP content of corneal nerves was positively correlated with 1 week OT score. There was also a significant increase in CD68-labeled macrophages in resveratrol-treated abraded corneas as compared to naïve corneas. These findings demonstrate the usefulness of our preclinical PRK model for the assessment of ocular pain therapeutics and indicate that topical resveratrol may not be useful for managing PRK-induced pain.

## Introduction

The cornea is a specialized multi-layered tissue that plays an essential role in vision and protects vulnerable ocular tissues from environmental insults. The outermost layer of the cornea is the corneal epithelium, which is densely innervated by sensory nerves. These corneal nerves are responsible for transducing and transmitting signals from potentially harmful environmental stimuli from the corneal epithelium via the trigeminal nerve to the trigeminal brainstem where the information is then conveyed to supraspinal substrates of nociception. Corneal sensory nerves are primarily nociceptive such that even low-threshold stimuli can evoke discomfort and pain. Corneal nerves are also part of the lacrimal functional unit, which maintains the tear film (Dartt and Willcox, 2013), a complex mixture of lipids, proteins, electrolytes and water that covers, nourishes and protects the ocular surface.

Vision correction surgeries, such as photorefractive keratectomy (PRK) and laser-assisted in situ keratomileusis (LASIK) were developed to take advantage of the accessibility of the cornea and its role in providing most of the focusing power of the eye, providing better vision to millions of patients. These procedures involve laser-induced ablation and reshaping of the corneal stroma, the thickest layer of the cornea that lies under the epithelium. The stroma is composed primarily of tightly packed collagen fibers that provide the transparency, structural integrity and curvature of the cornea, making it the main refractory component of the visual system (Sridhar, 2018). In order to access the stroma, a portion of the corneal epithelium must be removed. In PRK, alcohol is used to soften the corneal epithelium and then mechanical debridement is used to remove the epithelium. As a result, corneal sensory nerves are severed in the abraded area, causing acute pain, photophobia and alterations in tear production leading to sensations of dryness, burning, and grittiness in postoperative PRK patients (Beheshtnejad et al., 2015;Bower et al., 2015;Colin and Paquette, 2006;Gaeckle, 2021;Palochak et al., 2020;Quinto et al., 2008;Ripa et al., 2020;Shetty et al., 2019;Zarei-Ghanavati et al., 2019). There has been inconsistent and poor clinical management of post-operative PRK symptoms, usually with oral and topical non-steroidal anti-inflammatory drugs (NSAIDs), narcotics or corticosteroids (Javadi et al., 2008;O’Brart et al., 1994;Palochak et al., 2020;Ripa et al., 2020;Shetty et al., 2019). Preclinical models of PRK can assess new therapeutics for post-operative PRK pain and tear dysfunction.

Resveratrol is a natural polyphenol and phytoestrogen that is enriched in red grapes and wine (Tillu et al., 2012) among other food sources and plant species (Delmas et al., 2021) and has weak estrogen activity (Fremont, 2000;Thaung Zaw et al., 2020). Numerous studies have demonstrated the anti-oxidant, anti-inflammatory, anti-angiogenic, neuroprotective and cardioprotective effects of resveratrol (Delmas et al., 2021;Fremont, 2000), highlighting its enormous therapeutic potential in a variety of diseases, even ocular diseases. Several preclinical studies have demonstrated resveratrol’s potential as an effective analgesic in neuropathic, inflammatory and post-surgical pain states using a variety of dosing and formulations. Oral administration of resveratrol attenuated mechanical allodynia in male mice (Yin Y. et al., 2019) and male rats (Yang et al., 2016) after chronic constriction injury of the infraorbital nerve. Mechanical and thermal paw withdrawal latencies were increased by intraperitoneal injections of resveratrol in male rats with Paclitaxel-induced neuropathic pain (Li X. et al., 2019). Intrathecal resveratrol attenuated mechanical allodynia in male rats with a spared nerve injury (Wang Y. et al., 2020) and female rats in a bone cancer model (Hao et al., 2020). Studies have also demonstrated the analgesic benefit of local or topical application of resveratrol after cutaneous injury. Local injection (Tillu et al., 2012) or application of resveratrol formulated in a topical cream (Burton et al., 2017) is effective in attenuating mechanical allodynia in male mice in a model of plantar incisional pain, and does not interfere with wound healing. Pretreatment of the male rat hind paw with topical resveratrol-loaded niosomal hydrogel reduced paw edema in the hours after carrageenan-induced inflammation (Negi et al., 2017). There have also been studies on the potential therapeutic effects of resveratrol in ocular diseases, although they have not focused on its analgesic properties (Abengozar-Vela et al., 2019; Li M. et al., 2021;Marino et al., 2013;Shetty et al., 2020;Tsai et al., 2015) (see reviews (Delmas et al., 2021;Fremont, 2000)).

We have previously described a corneal abrasion model that simulates the initial step in the PRK procedure in which the corneal epithelium is removed and the corneal sensory nerves are severed in the abraded area (Hegarty et al., 2018). Furthermore, the behavioral, homeostatic and morphological effects of this corneal abrasion model are similar to the chief complaints of acute pain, photophobia and tear dysfunction from post-operative PRK patients. In light of these similarities, we used corneal abrasion in rats as a preclinical model to assess the ability of topical resveratrol formulated in ophthalmic ointment to reduce ocular pain and prevent decreases in tear production after PRK. In this study, we have also expanded our behavioral assessments to include orbital tightening, a component of the Rat Grimace Scale (Sotocinal et al., 2011), as a measure of spontaneous pain and possibly photophobia (Harris et al., 2017). Our study demonstrates the importance of a multi-faceted approach to measuring therapeutic effectiveness, thoughtful consideration of the therapeutic formulation, and the need to assess effectiveness of therapeutics in both sexes.

### Experimental Procedures Animals

Methods were approved by the OHSU Institutional Animal Care and Use Committee (IACUC). Experiments were performed on male and female Sprague-Dawley rats (8-9 weeks old; males: 250-290 g; females: 180-215 g); Charles River Laboratories, Wilmington, MA) that were housed in same sex pairs on a 12/12 light/dark cycle and had access to food and water at all times.

Experiments on males and females were conducted during different weeks so that rats from only one sex were in the testing room at a given time. A total of 38 rats (12 females, 26 males) were used with specific group sizes indicated in the Results section for each study. In the course of measuring behavioral responses, tear production, and corneal nerve density, some rats were assessed for more than one metric when possible.

### Corneal abrasions

Corneal abrasions were performed with heptanol as described previously (Hegarty et al., 2018). Briefly, rats were anesthetized with isoflurane and received topical proparacaine ophthalmic drops onto the surface of the left eye. A small metal ring (6-gauge (4.4 mm internal diameter) stainless steel tubing, custom, HTX-06R, Small parts, Inc, Miramar, FL) was secured to the left cornea with petroleum jelly, then heptanol (10 ul) was pipetted into the ring and left in place for 90 seconds. The heptanol was then wicked away, the corneal surface was rinsed with saline, and the damaged corneal epithelium was carefully debrided with a cotton swab. Subsets of rats received topical treatment of ophthalmic ointment with or without resveratrol immediately after debridement for 10 minutes while still anesthetized (see Figure 1). Rats awoke from anesthesia on a warming pad and were returned to their home cage and monitored until fully awake and mobile.

**Figure 1.**
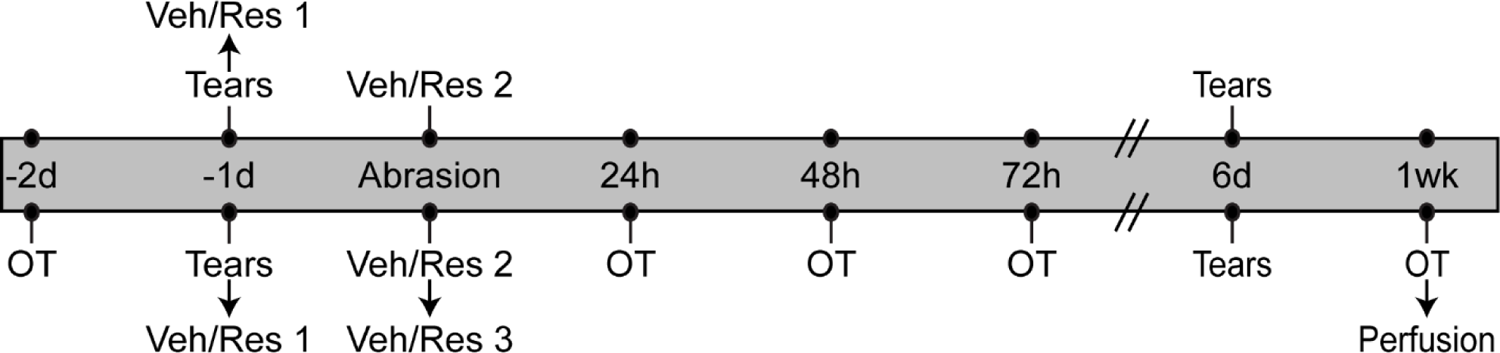
Timelines of assays and procedures for the resveratrol study. (**Top**) The tear production (Tears) and dosing schedule for our initial resveratrol dose-response study. (**Bottom**) The orbital tightening (OT) timeline includes OT behavior, tear production (Tears), dosing schedule with vehicle (Veh) or resveratrol (Res) and perfusion from pre-abrasion baseline time points (−2 days (d), −1d) through abrasion to acute post-abrasion time points (24h, 48h and 72h) and 1 week (1wk) post-abrasion. The dose number of vehicle or resveratrol is indicated (Veh/Res 1-3). Arrows denote the sequential order of assays and procedures that were done on the same day. There were no assays performed on days 4 and 5 (double lines).

### Phenol thread test

Tear production was assessed using the phenol thread test (Zone-Quick, Oasis Medical, Gendora, CA) as described previously (Aicher et al., 2015;Hegarty et al., 2018). This test is equivalent to the Schirmer’s test (Vashisht and Singh, 2011) used clinically to measure tear production. Rats were briefly anesthetized with isoflurane and placed on either their left or right side. One end of the phenol thread was placed in the lateral canthus of the eye for 15 seconds.

At the end of 15 seconds, the thread was removed and the length of thread that turned red was measured to the nearest millimeter (mm). Rats were then placed on their other side and the process was repeated. The eye order that tear production was measured was consistent for an individual animal throughout the study, however eye order was mixed among the rats within a treatment group. Each rat was tested one day prior to corneal abrasion (Baseline) and at 6 or 7 days (both labeled as 1 week) post-abrasion. Tear production tests were not performed on the same day as behavioral assessments as we have observed that even brief anesthesia interferes with behavioral testing on the same day.

### Orbital tightening

Spontaneous pain was assessed using orbital tightening (OT), which is one of the components of the Rat Grimace Scale (Sotocinal et al., 2011). Rats were recorded using two cameras (Hero 7 Silver and Hero 8 Black, Go Pro, San Mateo, CA, 1440 pixels, 30 frames/sec, Wide field of view (FOV)) in fixed positions on either side of an acrylic behavioral chamber (8.25 x 4.25 x 6.0” (L x W x H) with a black textured floor) in ambient light with the chamber cover on (approximately 500 lux; Urceri MT-912 light meter). Mirrors were positioned around the chamber to ensure continuous recording of the left eye. Rats were habituated to handling, the behavioral chamber, testing room, and cameras prior to the first OT assessment. On the testing day, rats acclimated to the testing room for at least 30 minutes prior to behavioral assessment. OT was assessed two days prior to corneal abrasion (Baseline) and then post-abrasion at 24, 48, 72 hours and 1 week (Figure 1). Rats were placed in the behavioral chamber to acclimate for 10 minutes, then the cameras recorded the rat for 5 minutes, then rats were returned to their home cage. Videos of each rat were converted to JPG images (Free Video to JPG Converter software, DVDVideoSoft). Ten images taken 30 seconds apart throughout the 5 minutes were extracted and then assessed for OT by two independent observers. Each image was scored for the degree of eye closure based on an ordinal scale described previously for rats (Sotocinal et al., 2011). Briefly, a score of “0” represented no eye closure or orbital tightening; a score of “1” represented a moderate amount of orbital tightening visible, with no more than one-half of the eye covered; and a score of “2” representing a substantial amount of orbital tightening with more than one-half of the eye covered, including complete eye closure. Efforts were made to ensure that images were not taken during a brief eye closure (i.e., blinking) or during grooming. Ten scores were collected for each rat and summed for that testing day.

### Immunohistochemistry

One week after corneal abrasion and after final behavioral assessments, rats were overdosed with sodium pentobarbital (150 mg/kg) and perfused transcardially through the ascending aorta with 10 ml of heparinized saline (1000 units/ml) followed by 600 ml of 4% paraformaldehyde in 0.1 M phosphate buffer, pH 7.4 (PB) (Hegarty et al., 2018;Hegarty et al., 2017). The eyes were enucleated immediately after perfusion and placed in PB. Using a 5 mm corneal trephine (Ambler Surgical, Exton, PA), the central corneas were removed and placed in fresh PB at 4 °C until immunoprocessing.

Whole mount corneas were immunoprocessed free-floating as described previously (Hegarty et al., 2018;Hegarty et al., 2017). Corneas were incubated for 3 nights at 4°C in a primary antibody cocktail of Mouse anti-β-tubulin (1:500, RRID: AB_10063408, Cat. 801202, Biolegend, San Diego, CA), Sheep anti-Calcitonin gene-related peptide (CGRP, 1:3000, RRID: AB_725809, Cat. ab22560, Abcam, Cambridge, MA) and Rabbit anti-CD68 (1 µg/µl, RRID: AB_2746112, Cat. PA5-78996, Invitrogen/ThermoFisher Scientific (TFS), Waltham, MA) in 0.25% Triton X-100 (Sigma-Aldrich, St. Louis, MO) and 0.1% Bovine Serum Albumin (BSA, Sigma) in 0.1 M Tris-buffered saline pH 7.6 (TS). Corneas were then rinsed and incubated in a fluorescent secondary antibody cocktail of Donkey anti-Rabbit AlexaFluor (AF) 488 (1:800, RRID: AB_2535792, Cat.

A-21206, TFS), Donkey anti-Mouse Cy3 (1:800, RRID: AB_2315777, Cat. 715-165-151, Jackson ImmunoResearch, West Grove, PA) and Donkey anti-Sheep AF647 (1:800, RRID: AB_2535865, Cat. A-21448, TFS) in 0.1% BSA in TS light-protected for 2 hours at room temperature. Corneas were rinsed and either blotted dry and placed into individual wells of an 18 well µ-slide (ibidi USA, Inc., Fitchburg, WI) and coverslipped with CFM-1 (Electron Microscopy Sciences, Hatfield, PA) or cloverleafed, flattened, and mounted onto gelatin-coated slides and coverslipped with ProLong Gold Antifade Mountant (TFS).

### Imaging and Image analysis

Antibodies to β-tubulin, CGRP, and CD68 were used to assess corneal nerve density, CGRP content within corneal nerves, and macrophage infiltration into the corneal epithelium, respectively, using methods similar to our previously described analyses (Hegarty et al., 2018;Hegarty et al., 2017). Confocal images were taken at the apex of the central cornea and were captured on a Zeiss LSM900 confocal microscope with a Plan-Apochromat 20x / 0.8 M27 objective (Carl Zeiss MicroImaging, Thornwood, NY) using the single pass, multi-track format at a 2048 x 2048 pixel resolution in the Advanced Light Microscopy Core (ALMC) at OHSU. Optical sectioning produced Z-stacks of 0.58 µm optical slices that were bounded by the extent of fluorescent β tubulin, CGRP, and CD68 immunolabeling throughout the thickness of the corneal epithelium.

Volumetric assessments of β tubulin-labeled cornea nerve density, CGRP content within the nerves, and CD68 in the epithelium were performed using Imaris 9.5 software (Bitplane USA, Concord, MA) on an offline workstation in the ALMC at OHSU by a treatment-masked observer as previously described in detail (Hegarty et al., 2018;Hegarty et al., 2017). Only the left cornea was assessed for each animal. Briefly, we defined the corneal epithelium by the β tubulin-and CGRP-labeled sub-basal and intraepithelial corneal nerves and background labeling of CD68 from the anterior surface of the cornea to the epithelial-stromal border. The corneal epithelium was isolated as a region of interest (ROI) using the Surfaces Segmentation tool and its volume was calculated by the software (µm^3^). The Mask Channel function was then used to isolate β-tubulin, CGRP, and CD68 labeling within the corneal epithelium ROI for further analysis. β tubulin labeling within the ROI was determined using the Thresholding function and Surfaces Segmentation tool in Imaris, and a volume (µm^3^) was calculated. In order to account for subtle differences in the corneal epithelium volumes, the volume of β tubulin labeling was calculated as a percentage of the corneal epithelium volume for each cornea and expressed as the mean % β tubulin or the percent of the epithelium that contains β tubulin-labeled nerves. To measure the CGRP content specifically within the corneal nerves, we established the β tubulin volume as the ROI and isolated the CGRP labeling using the Mask Channel function and then CGRP volume (µm^3^) was calculated. The volume of corneal nerve CGRP labeling was calculated as a percentage of the epithelial volume and expressed as % CGRP. CD68 labeling was also measured using the Surfaces Segmentation Tool and the CD68 volume (µm^3^) was calculated as a percentage of epithelial volume and expressed as % CD68.

### Statistical analysis

All statistical analysis was performed using SigmaPlot 12.0 software (Systat, Palo Alto, CA). Two-way repeated measures ANOVA were used to compare OT scores, and Holm-Sidak all pairwise multiple comparison post hoc tests were performed when there was a statistically significant difference in one or more factors or the interaction of those factors. We performed paired t-tests for within-animal comparisons of raw phenol thread measurements taken at baseline and 1 week except in the case when the normality test failed and the non-parametric Wilcoxon Signed Rank test was performed instead. Two-way ANOVAs with Holm-Sidak all pairwise multiple comparison post hoc tests were also performed to compare measurements of corneal epithelium volume, % β tubulin, % CGRP and % CD68 in female and male treatment groups. Linear regressions were performed to examine the relationship between OT behavior and tear production, % CGRP or % CD68.

### Timeline of assessments and drug applications

All animals undergoing orbital tightening assessment received an abrasion to the left cornea and received three applications of drug to the left eye (Figure 1, bottom timeline). Orbital tightening (OT) assessment was performed two days prior to corneal abrasion (−2d) and at several post-abrasion time points (24, 48, 72 hours and 1 week; Figure 1). A separate group of abraded male rats were assessed for tear production as part of an initial dose-response study using 0% (vehicle), 1%, 2% or 4% resveratrol (Figure 1, top timeline). These animals only received two doses of drug at −1d (Veh/Res 1) or immediately after Abrasion (Veh/Res 2). All assays were performed on the left (abraded) eye.

The pre-abrasion drug application was performed 1 day before corneal abrasion in anesthetized animals, immediately after baseline tears were measured (−1d; Figure 1). Two drops of proparacaine were added to the left cornea, then the metal ring was secured to the ocular surface with petroleum jelly. Vehicle or resveratrol was added to the ring with a syringe such that the ocular surface within the ring was completely covered (∼0.03 ml) for 10 minutes. The ring was removed, but the drug was allowed to remain on the ocular surface as the animals recovered on the heating pad and in the home cage. Drug was also applied immediately after abrasion (Veh/Res 2; Figure 1) on anesthetized animals using the same procedure described above. For animals in the OT group, there was also one more drug application 3-4 hours post-abrasion when the rats were awake (Veh/Res 3; Figure 1). For these applications, each rat was held in a soft towel and drug was gently applied to the left eye with a cotton-tipped applicator.

Each rat was then placed into the behavior chamber for 10 minutes before returning to the home cage to avoid transferring topical ointment to a cage mate. Tear production was assessed on day 6 or 7 after corneal abrasion. After the final OT assessment at 1 week, rats were perfused with aldehydes for histological studies.

### Resveratrol Ophthalmic ointment formulations and preparation

The OHSU Medicinal Chemistry Core worked on various formulations of resveratrol (*trans*-resveratrol, CAS No. 501-36-0, product no. 70675, Cayman Chemical, Ann Arbor, MI) to find a delivery method that would be compatible with topical application to the eye. Resveratrol is quite difficult to dissolve without the use of chemicals that would be incompatible with administration into the eye and therefore we settled on a formulation that was a suspension in an emulsion ointment. The chemists provided 3 different doses of resveratrol (1%, 2%, 4%) and vehicle ointment (0%) to be tested and the experimenter was blinded to the contents during all assessments.

To prepare the ophthalmic formulations, all equipment was wiped with 70% ethanol and preparation was done using sterile methods next to a Bunsen burner flame. The appropriate amount of resveratrol was weighed into a flat-bottomed glass cover and the corresponding amount of Sodium Chloride Ophthalmic Ointment USP, 5 % (in mineral oil, modified lanolin, purified water and white petrolatum; Akorn Pharmaceuticals, Lake Forest, IL) was added. The suspension was homogenized using two Corning Costar polyethylene cell lifters to spread out and mix the two components until they formed a homogeneous paste (no streaks or clumps when spreading). The paste was then transferred into a sterile 1 mL HENKE-JECT Tuberculin syringe (Air-Tite Products Co., Inc., Virginia Beach, VA) and sealed with a syringe cap. Uniform suspension in the emulsion was verified by the Medicinal Chemistry Core on a Zeiss Telaval 31 inverted microscope and visualized on an Olympus BX51 microscope using a 20x / 0.50 UPlanFL N objective equipped with a DP74 camera and associated cellSens software (Olympus America, Center Valley, PA) (Figure 2). Images used for publication were converted to grayscale and adjusted for optimal brightness and contrast using Adobe PhotoShop CS5 (Adobe, San Jose, CA).

**Figure 2.**
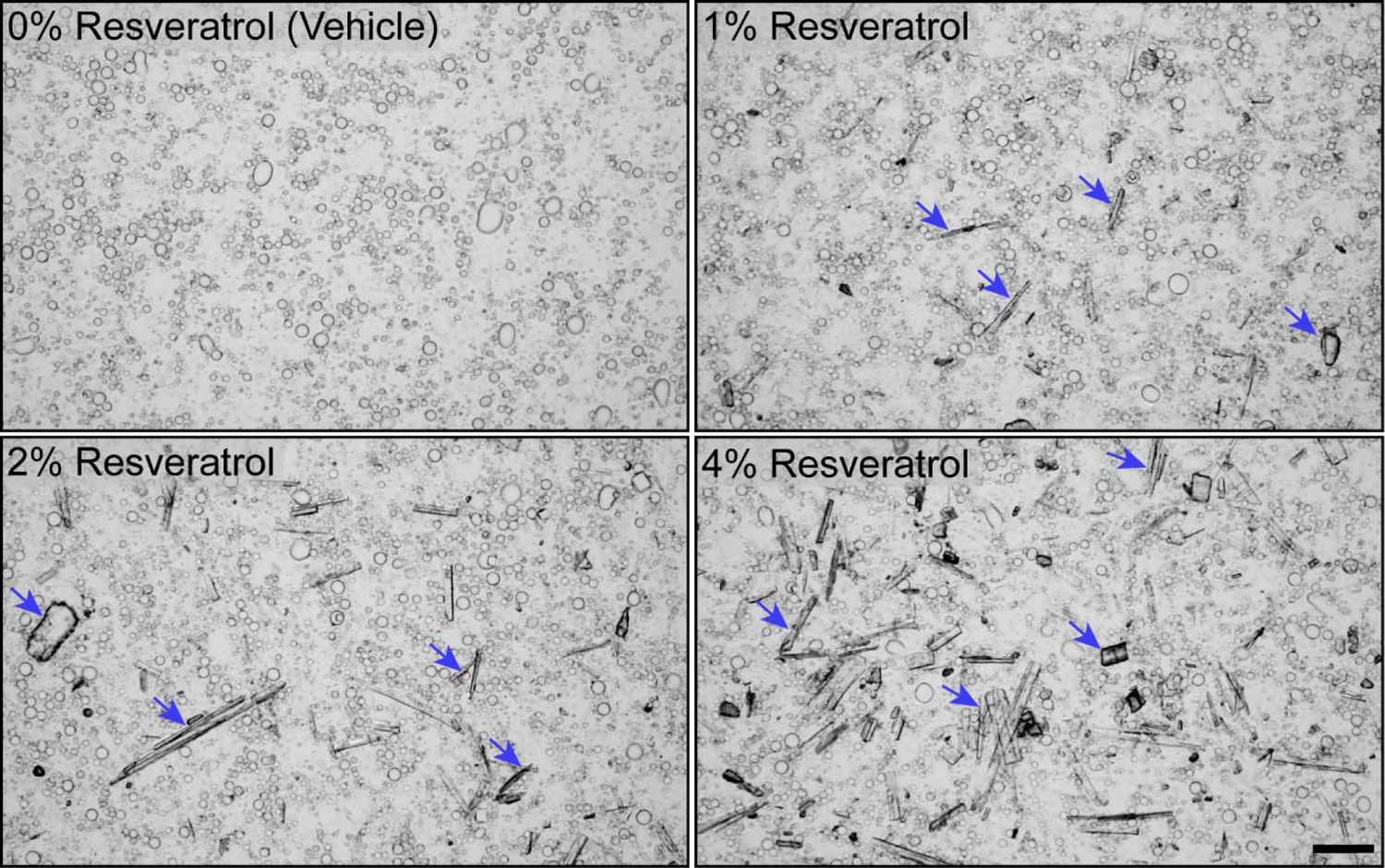
Suspensions of resveratrol in ophthalmic ointment used for topical ocular application. Resveratrol crystals are rod-like structures of various sizes in suspension (blue arrows). Scale bar = 50 µm.

## Results

### Topical application of resveratrol increases tear production after corneal abrasion

Tear production was measured in abraded male rats that received two applications of resveratrol (Figure 1, Veh/Res 1 and 2) as part of our initial dose-response study (0% (vehicle), 1%, 2%, 4%). Some animals in the vehicle and 4% resveratrol groups were also assessed for OT behaviors. We used paired t-tests to make within-animal comparisons in each group (Table 2). Male rats that received 2% or 4% resveratrol demonstrated a significant increase in tear production 1 week post-abrasion compared to Baseline values (Table 2). There were no changes in tear production in the unabraded right eye (data not shown). Our previous studies in abraded, untreated animals (Hegarty et al., 2018) showed a reduction in tear production at 1 week after corneal abrasion, which we did not see in the present study. These findings suggest that the ophthalmic ointment itself may be preventing or masking the normal decrease we usually see at 1 week post-abrasion. We only tested vehicle and 4% resveratrol in females and we found no change in tear production in females in either treatment group (Table 2).

**Table 1.**
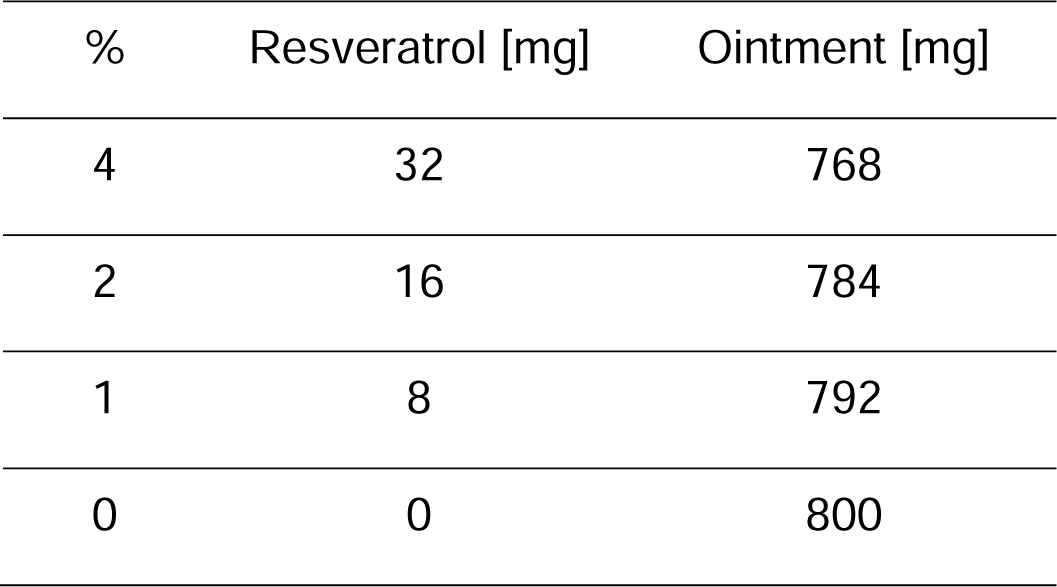
Concentrations of resveratrol in ophthalmic ointment

**Table 2.** Raw phenol thread (PT) measurements taken from the abraded eye before (Baseline) and 1 week after abrasion.

| Sex | Treatment | Baseline PT | 1 week PT | P value | N |
| --- | --- | --- | --- | --- | --- |
| Male | Vehicle | 7.3 ± 1.3 mm | 8.2 ± 0.7 mm | 0.210 | 9 |
| Male | 1% resveratrol | 11.3 ± 3.2 mm | 11.0 ± 3.0 mm | 1.000 | 3 |
| Male | 2% resveratrol | 8.7 ± 1.8 mm | 15.3 ± 1.9 mm * | 0.00853 | 3 |
| Male | 4% resveratrol | 7.4 ± 1.0 mm | 12.4 ± 2.7 mm * | 0.0402 | 7 |
| Female | Vehicle | 14.5 ± 1.4 mm | 14.3 ± 2.7 mm | 0.465 | 4 |
| Female | 4% resveratrol | 10.8 ± 1.8 mm | 12.5 ± 2.1 mm | 0.201 | 4 |

PT measurements are expressed as mean ± SEM mm. Male vehicle and 4% resveratrol groups include rats that received 2 doses of drug from the dose-response study and 3 doses from the OT behavioral study. Paired t-tests were used for most comparisons. A Wilcoxon Signed Rank test was used for the male 1% resveratrol group. \* P < 0.05 vs Baseline value

### Corneal abrasion increases spontaneous pain in male and female rats

We used the orbital tightening (OT) metric of the Rat Grimace Scale (Sotocinal et al., 2011) to measure spontaneous pain in female and male rats before and at several time points after corneal abrasion (Figure 3). Two different control groups received either no treatment (naïve) or an ointment that did not contain active drug (vehicle). For each sex, we compared the OT scores of naïve and vehicle groups using a two-way repeated measures ANOVA. We found no significant differences between these groups, so results were pooled for each sex and are referred to as control. We found that following corneal abrasion, OT scores were increased at 24, 48 and 72 hours and 1 week after abrasion compared to Baseline in both male and female control rats (Figure 3, two-way ANOVA, P < 0.001 for all time points). OT scores at 24 and 48 hours were significantly greater than those measured at 72 hours (P < 0.001; P = 0.003, respectively). At 24, 48 and 72 hours OT scores were all significantly greater than those at 1 week (P < 0.001 for all comparisons). There was no effect of sex (P = 0.410) and no significant interaction between time point and sex (P = 0.154). These data demonstrate that females and males have a similar OT time course after corneal abrasion, with responses elevated acutely, decreasing by 72 hours and approaching baseline levels at 1 week, suggesting a significant but incomplete recovery to pre-abrasion baseline behavior.

**Figure 3.**
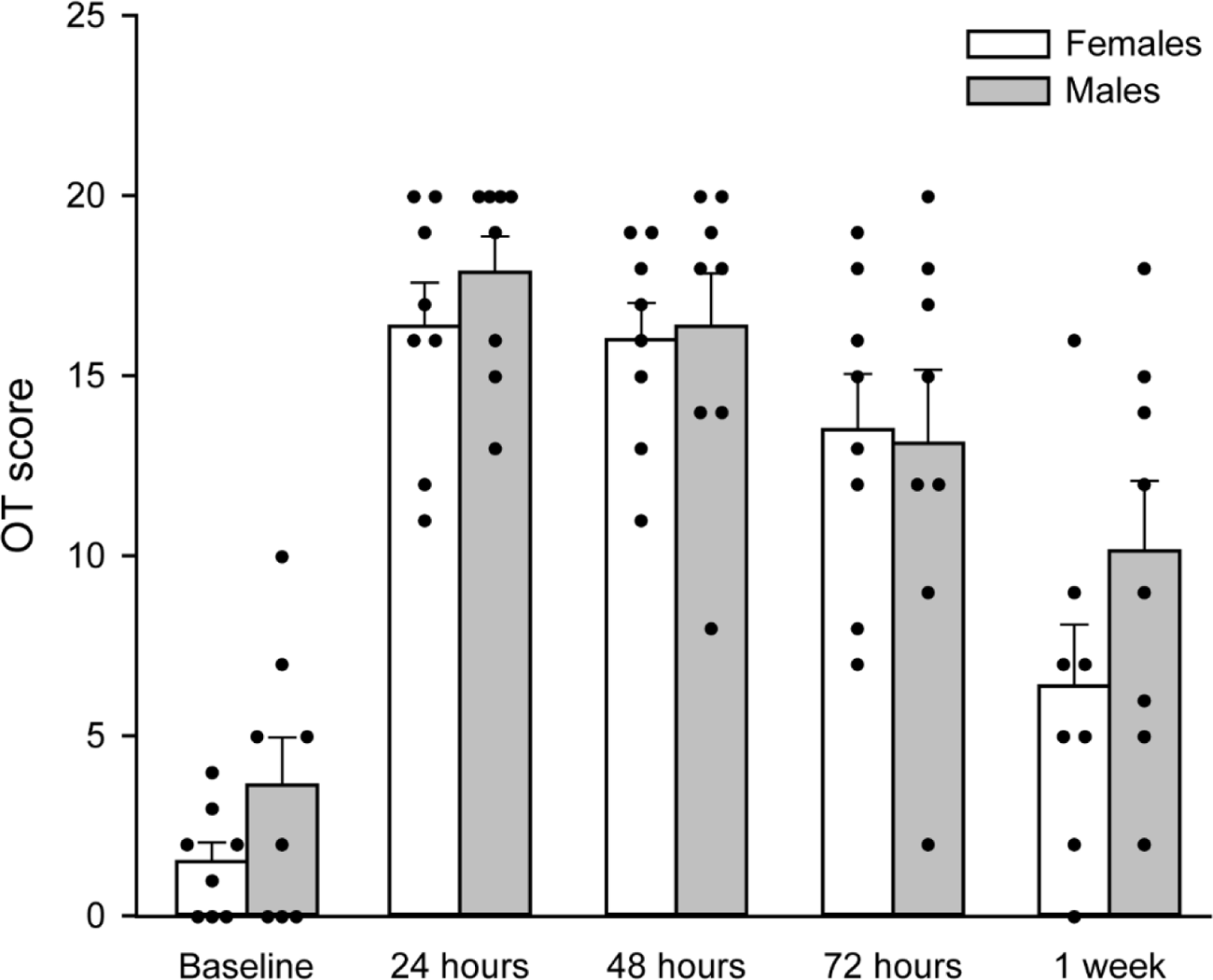
Female and male control rats show a similar time course of orbital tightening (OT) behavior after corneal abrasion. Individual rats for each group are represented by dots. White (Females) and gray (Males) bars represent mean OT scores ± SEM. n = 8 females; 8 males.

### Spontaneous pain is prolonged after 4% resveratrol in males

We compared OT scores for female and male rats that received topical application of 4% resveratrol to the ocular surface after corneal abrasion (Figure 4). A two-way repeated measures ANOVA found a statistically significant interaction between sex and time point (P = 0.006). Post-hoc comparisons found significant increases in OT scores in resveratrol females at all post-abrasion time points compared to Baseline (24 and 48 hours, P < 0.001; 72 hours, P = 0.002; 1 week, P = 0.031). In addition, OT scores at 1 week were reduced compared to 24 and 48 hours (P < 0.001), but not 72 hours (P = 0.317). The 72 hour OT score was significantly less than the 24 hour (P = 0.002) and 48 hour (P = 0.019) OT scores. In comparison, resveratrol males experienced significant increases in OT scores at all post-abrasion time points (P < 0.001 for all time point vs. Baseline). When comparing males and females, there was a significant difference between the two sexes at 72 hours and 1 week (Figure 4, black lines, P < 0.001 for both time points). Within-sex two way repeated measures ANOVAs demonstrate that there are no significant differences between control females (Figure 3) and resveratrol females (Figure 4). Similar comparisons of control males (Figure 3) to resveratrol males (Figure 4) confirm significant differences in OT Scores at 72 hours and 1 week (P = 0.037 for both time points) for males only. These data suggest that topical ocular resveratrol application offers no benefit to females or males and may have a detrimental effect in males after corneal abrasion.

**Figure 4.**
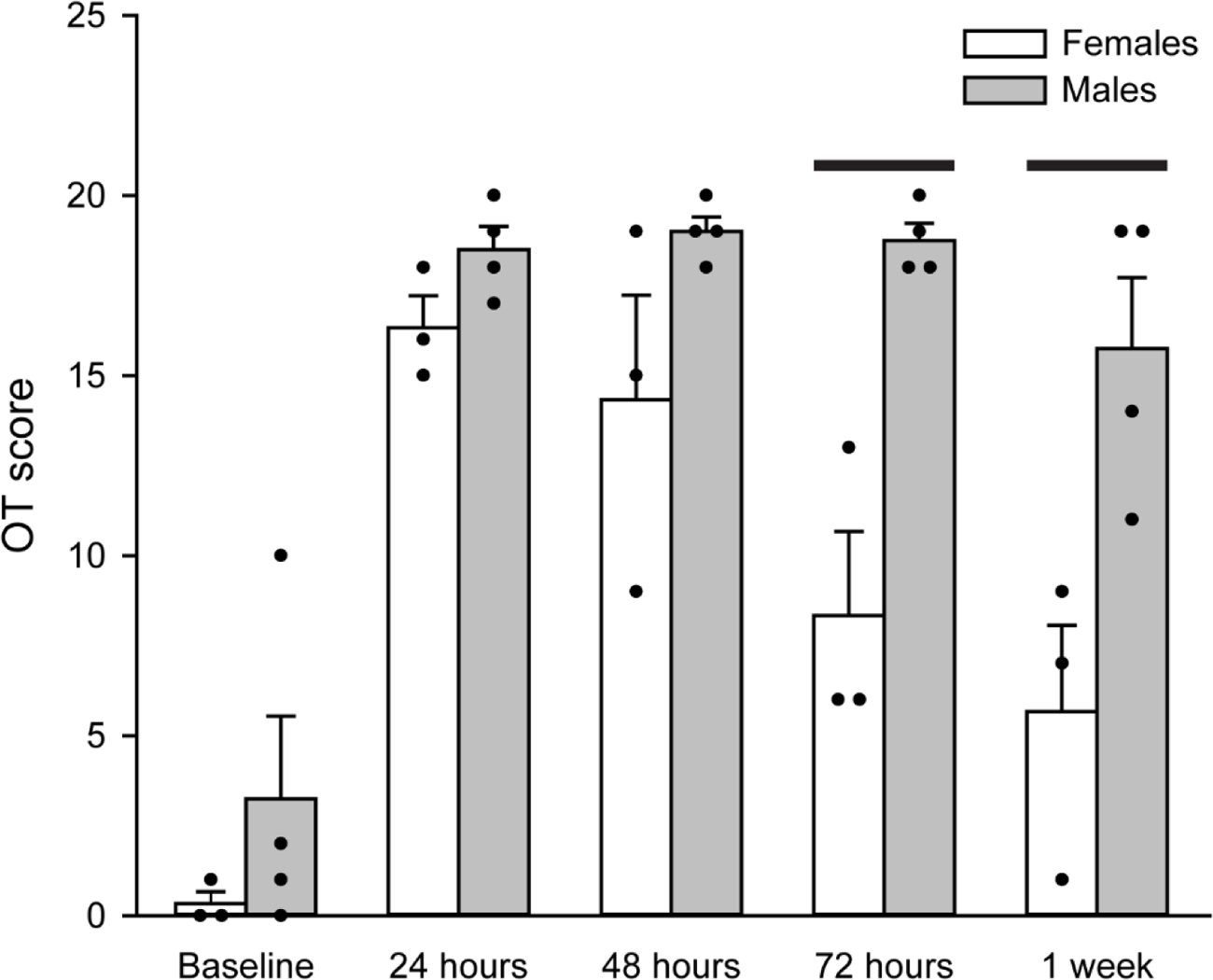
Resveratrol has different effects on spontaneous pain in male and female rats after corneal abrasion. Individual rats for each group are represented by dots and white (Females) and gray (Males) bars represent mean OT scores ± SEM. n = 3 females; 4 males. Black lines: P < 0.001.

We noted that there was individual variance in both tear measurements and OT scores (Table 2; Figures 3 - 4), so we assessed whether OT scores were correlated with tear measurements at 1 week. There is no relationship between tear production and spontaneous pain behavior (Figure 5, black line = linear regression, n = 15, R = 0.327, R^2^ = 0.107, Adjusted R^2^ = 0.0384, P = 0.234).

**Figure 5.**
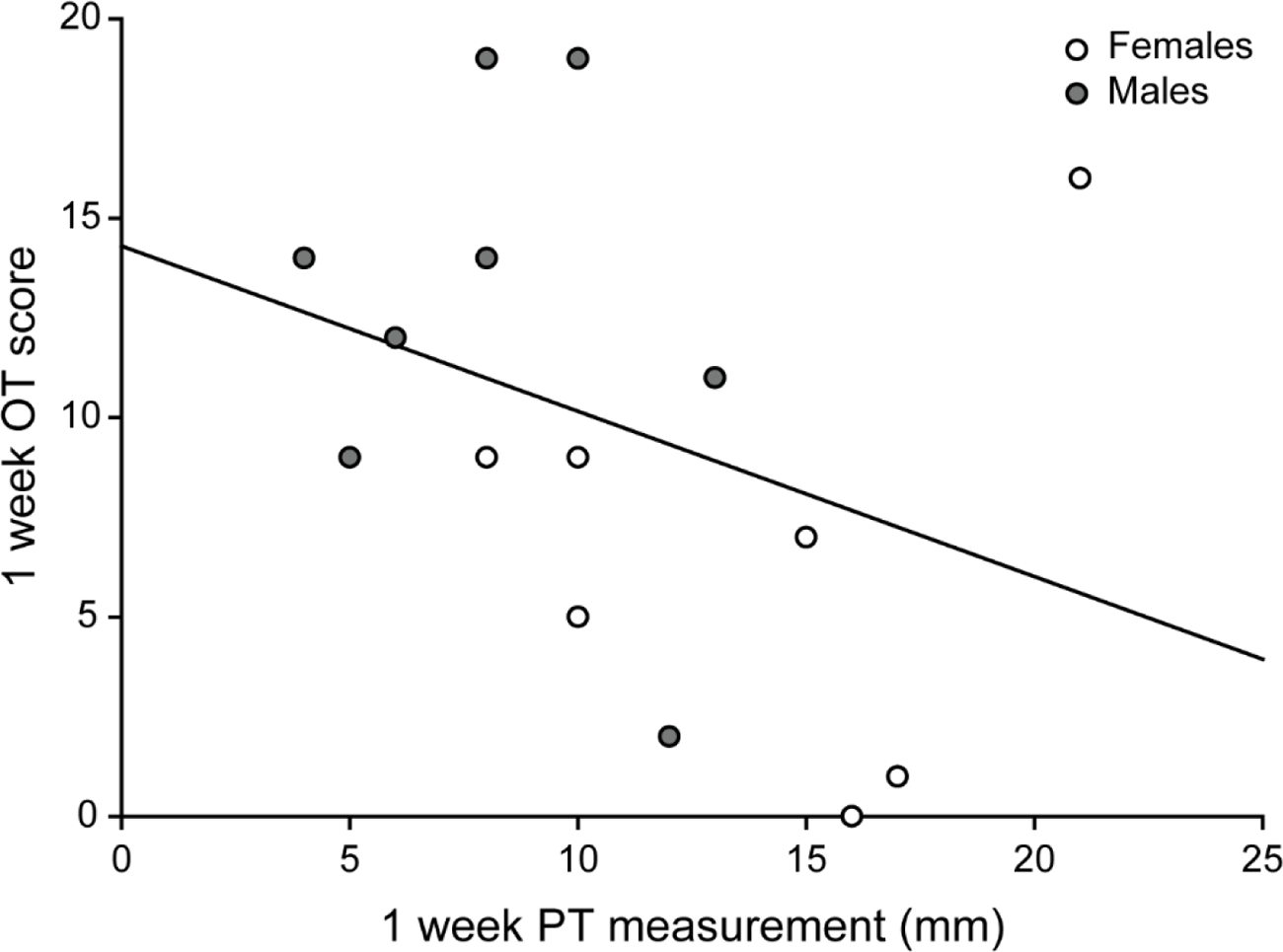
There is not a significant correlation between spontaneous pain (1 week OT score) and tear production (1 week PT measurement (mm)) that would account for the variance observed in the raw PT (phenol thread) measurements and OT scores.

### Corneal epithelial volume and corneal nerve density are reduced by corneal abrasion

The corneal epithelium normally heals quickly after an abrasion, and pain then resolves within one week in the rat (and most humans). However, failure of the epithelium to heal can produce ongoing pain. Therefore, we assessed corneal epithelial thickness, corneal nerve density, CGRP content and CD68 infiltration in rats treated with 4% resveratrol (n = 6, 3 females, 3 males) or vehicle (n = 8, 4 females, 4 males) one week after abrasion to assess the efficiency of epithelial restoration (Figure 6). A group of naive females (n = 3) and naive males (n = 4) who did not receive a corneal abrasion or any treatment were used as controls. We found a significant difference in epithelium volume (Figure 7A) among treatment groups (two-way ANOVA, P < 0.001) but no differences between females and males (P = 0.303) or interactions between treatment group and sex (P = 0.940). Corneal epithelium volumes were significantly decreased in abrasion animals that received 4% resveratrol (P < 0.001) or vehicle (P = 0.002) as compared to naïve animals. There was no significant difference in epithelium volume between the abrasion groups (resveratrol vs vehicle, P = 0.279). β-tubulin-labeled corneal nerve density was also analyzed among the treatment groups at 1 week post-abrasion (Figure 6, 7B). Similar to epithelium volume, we found a significant difference in corneal nerve density among treatment groups (two-way ANOVA, P < 0.001), but no differences between females and males (P = 0.490) or interactions between treatment group and sex (P = 0.099). Corneal nerve density was significantly reduced in abrasion animals that were treated with 4% resveratrol (P < 0.001) or vehicle (P = 0.014) as compared to naive animals. There was no significant difference in corneal nerve density between the abrasion groups (resveratrol vs vehicle, P = 0.137). These data demonstrate that ocular resveratrol treatment does not accelerate recovery of the corneal epithelium or corneal nerve density after abrasion.

**Figure 6.**
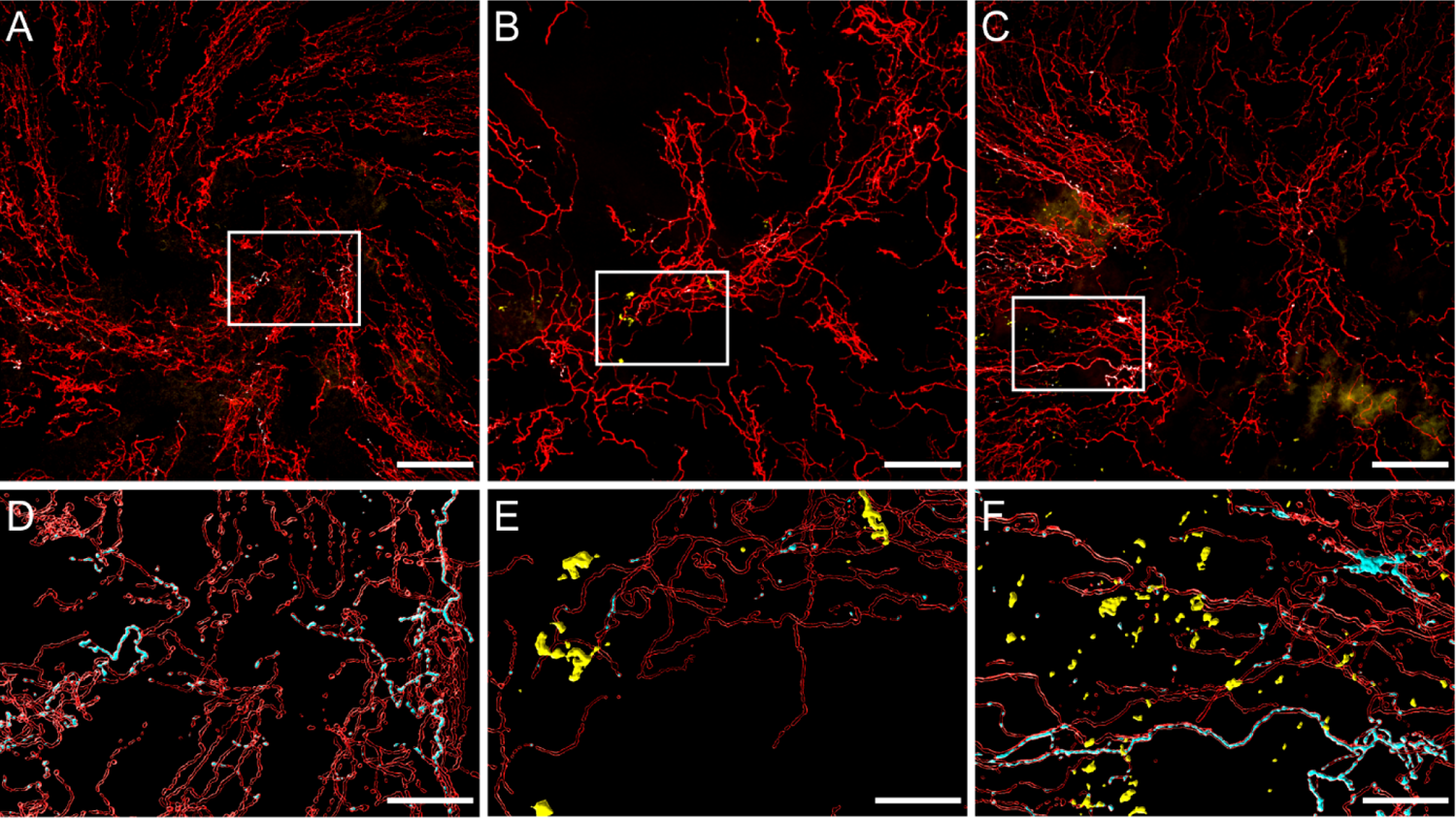
Corneal nerve density, CGRP content and macrophage infiltration were measured in corneas from naïve and abraded rats that received vehicle or 4% resveratrol. (**A – C**) Representative confocal micrographs of β tubulin-labeled corneal nerves (red), CGRP-labeled nerves (cyan) and CD68-labeled macrophages (yellow) in corneas from a naïve female (**A, D**), an abraded resveratrol female (**B, E**) and an abraded resveratrol male (**C, F**). The regions within white boxes are represented at higher magnification in panels D – F as volumetric renderings of all three labels. The renderings of β tubulin-labeled corneal nerves (red) were made transparent in order to see the CGRP renderings (cyan) within the corneal nerves. CD68-labeled macrophages (yellow) were more abundant in the abraded resveratrol male (**C, F**) compared to the abraded resveratrol female (**B, E**) and none were present in the naïve female cornea (**A, D**). Panels A – C scale bar = 100 µm; Panels D - F scale bar = 30 µm.

**Figure 7.**
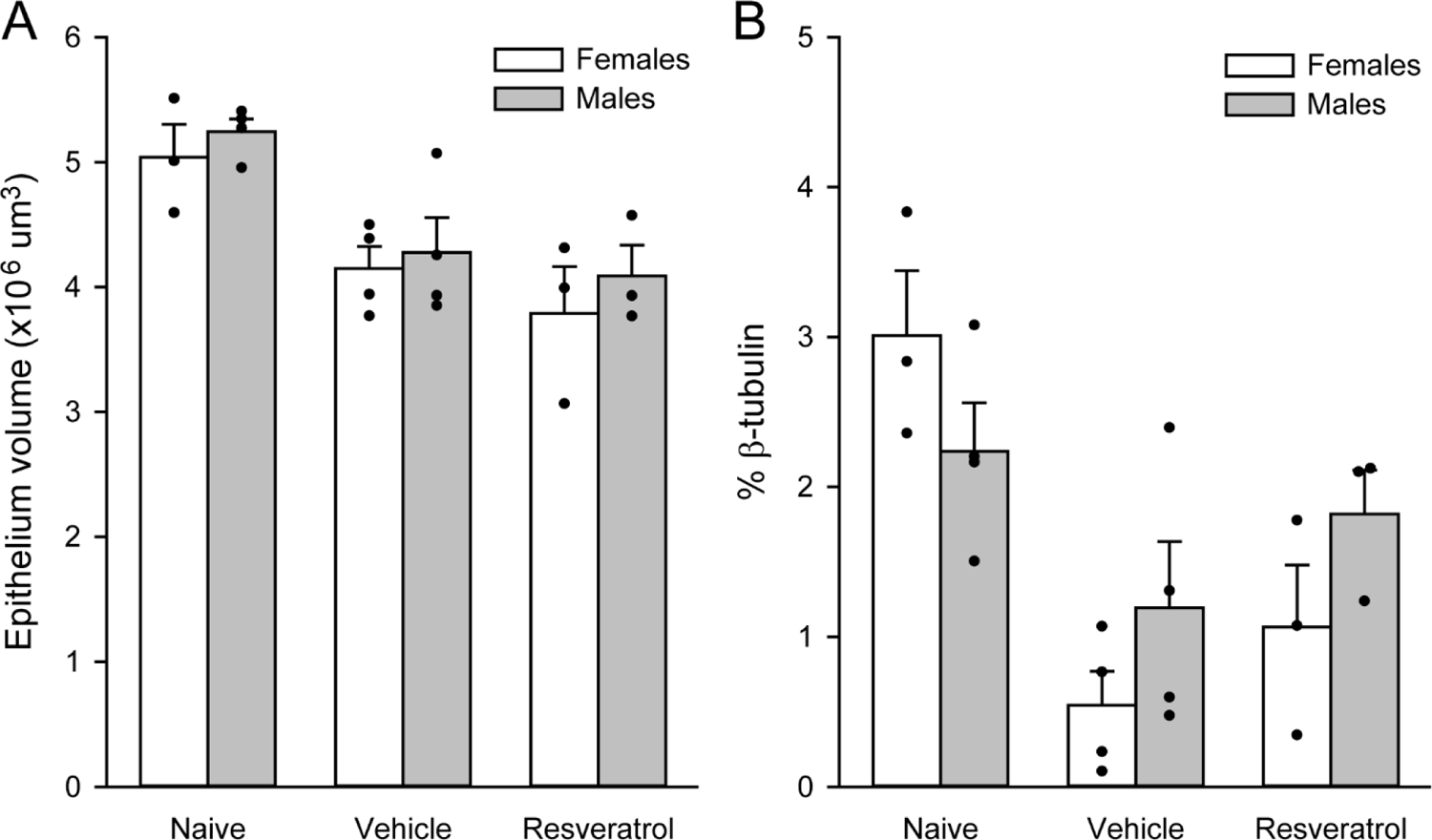
Corneal epithelium volume and corneal nerve density are decreased after abrasion and are not affected by resveratrol treatment. (**A**) Epithelial volume and (**B**) corneal nerve density, measured as the percent of the volume of β tubulin-labeled nerves within the epithelium volume (% β tubulin), were significantly decreased in rats that received a corneal abrasion as compared to naïve rats, regardless of treatment (vehicle, resveratrol). Individual rats for each group are represented by dots and bars represent mean epithelium volume or mean % β tubulin ± SEM.

### CGRP content in corneal nerves is decreased in females after corneal abrasion

CGRP content of corneal nerves was analyzed among the treatment groups at 1 week post-abrasion (Figure 6, 8). The % CGRP data was rank transformed in order to meet normality and equal variance criteria for a two-way ANOVA. There was a significant interaction between treatment group and sex (two-way ANOVA, P = 0.036). Post hoc comparisons demonstrated a significant difference between abraded females and abraded males that received resveratrol (Figure 9, black line; P = 0.014). Among female treatment groups (Figure 8, white bars), there was a significant reduction in % CGRP in abraded animals that received resveratrol (P = 0.010) and vehicle (P = 0.021) as compared to naïve animals. There were no significant differences among the male treatment groups (Figure 8, gray bars).

**Figure 8.**
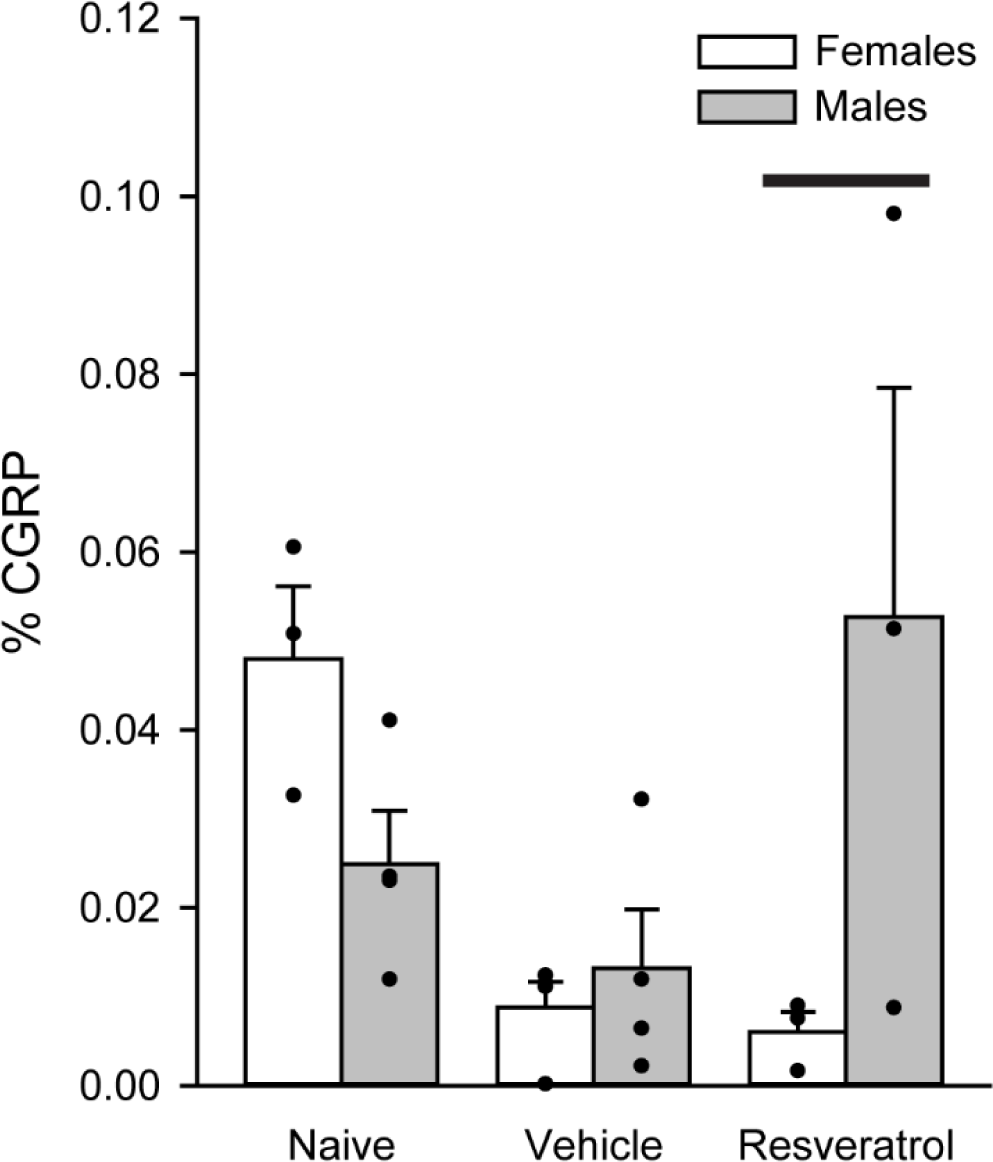
The percent of the corneal epithelium that contains CGRP-labeled nerves (% CGRP) is different in female and male resveratrol-treated rats. The % CGRP in female abraded corneas that received resveratrol was significantly less than the % CGRP in abraded male corneas that also received resveratrol (black line). Within the female treatment groups, % CGRP was decreased in abraded corneas as compared to naïve corneas, regardless of treatment (vehicle, resveratrol), while in males, there were no significant differences among any treatment groups. Non-transformed % CGRP data from individual rats for each group are represented by dots and bars represent mean % CGRP ± SEM.

**Figure 9.**
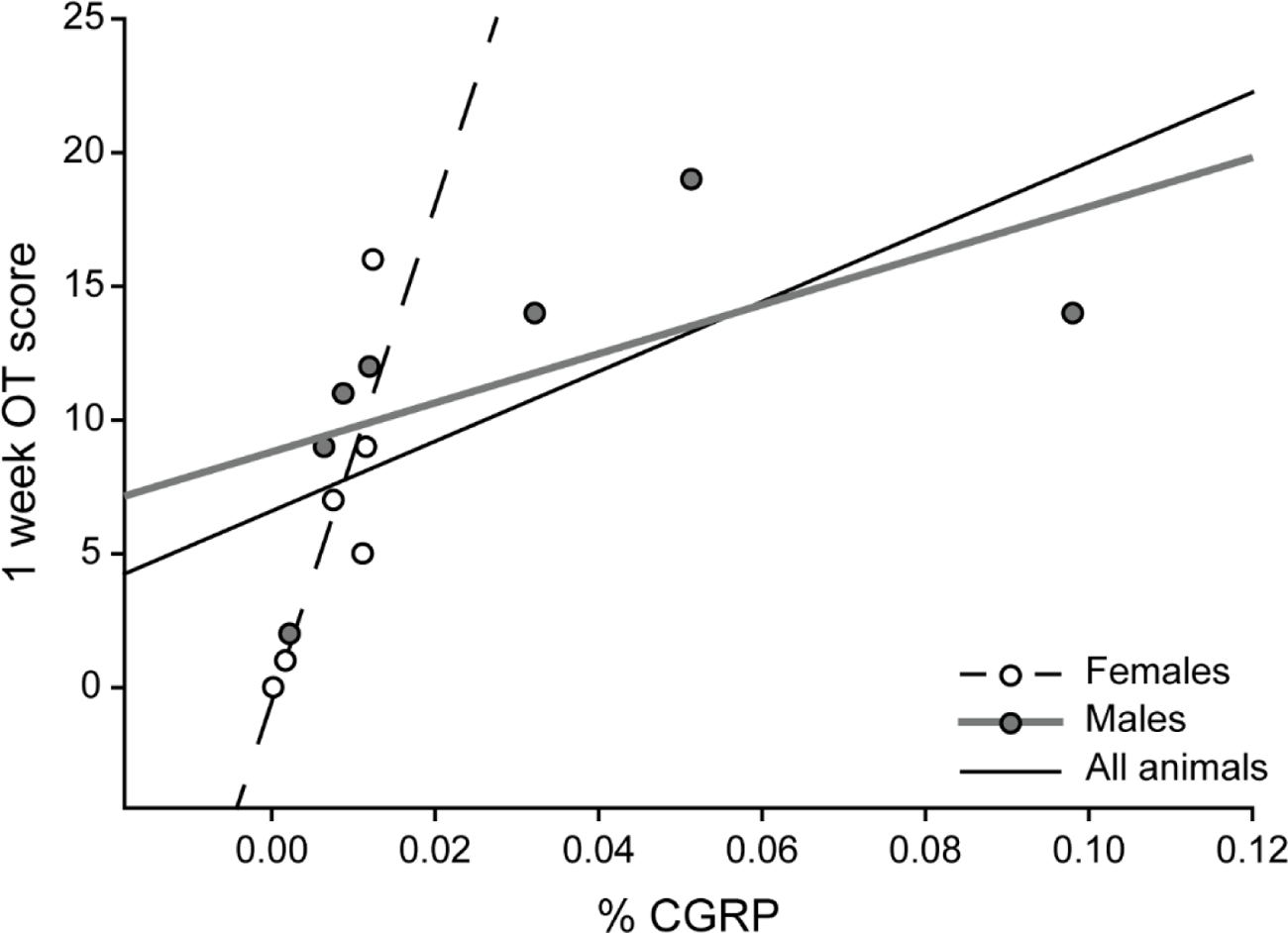
There is a significant correlation between spontaneous pain (1 week OT score) and corneal % CGRP (solid black line) for all animals, but this relationship appears to be driven by female rats (white dots, dotted line), rather than males (gray dots, gray line). % CGRP values from individual rats were used for linear regression analyses.

We assessed whether OT scores were correlated with corneal % CGRP. When all abraded rats were analyzed together, there was a significant relationship between the OT score at 1 week and the density of CGRP labeling in the corneal nerves (Figure 9, black line = linear regression, n = 6 females, 7 males; R = 0.600, R^2^ = 0.360, Adjusted R^2^ = 0.302, P = 0.03). When females and males were analyzed separately, OT score at 1 week and % CGRP in females were correlated (Figure 9, dotted line = linear regression, n = 6 females; R = 0.841, R^2^ = 0.708, Adjusted R^2^ = 0.635, P = 0.036), with small changes in corneal nerve CGRP related to large differences in OT behavior at 1 week post-abrasion. There was no relationship between % CGRP and OT score in male rats (Figure 9, gray line = linear regression, n = 7 males; R = 0.605, R^2^ = 0.366, Adjusted R^2^ = 0.239, P = 0.150). These data demonstrate that the relationship between CGRP and spontaneous pain is more tightly linked in females than males after corneal abrasion, consistent with other recent studies (Paige et al., 2022).

### CD68 labeling in the cornea is increased after resveratrol treatment

The volume of CD68 labeling was measured and normalized to the corneal epithelium volume (% CD68) and compared across treatment groups at 1 week post-abrasion (Figure 10). The % CD68 data was rank transformed in order to meet normality and equal variance criteria for a two-way ANOVA. There was a significant difference among treatment groups (P = 0.016) but no effect of sex (P = 0.170) and no interaction between treatment group and sex (P = 0.879). Post hoc comparisons found that resveratrol-treated abraded corneas had a significantly higher %CD68 as compared to naïve corneas (Figure 10; P = 0.015). The % CD68 in vehicle-treated abraded corneas was not significantly different from either naïve corneas (P = 0.281) or resveratrol-treated corneas, although there was a trend (P = 0.069). These data demonstrate that resveratrol increases macrophage infiltration into abraded corneas, although the effect of corneal abrasion alone cannot be completely ruled out. We also assessed whether there was a relationship between % CD68 and OT score at 1 week that could account for the variance in the data, especially in the male rats. There was no relationship between % CD68 in the cornea and spontaneous pain behavior at 1 week (Linear regression, n = 13, R = 0.263, R^2^ = 0.0692, Adjusted R^2^ = 0.000, P = 0.385).

**Figure 10.**
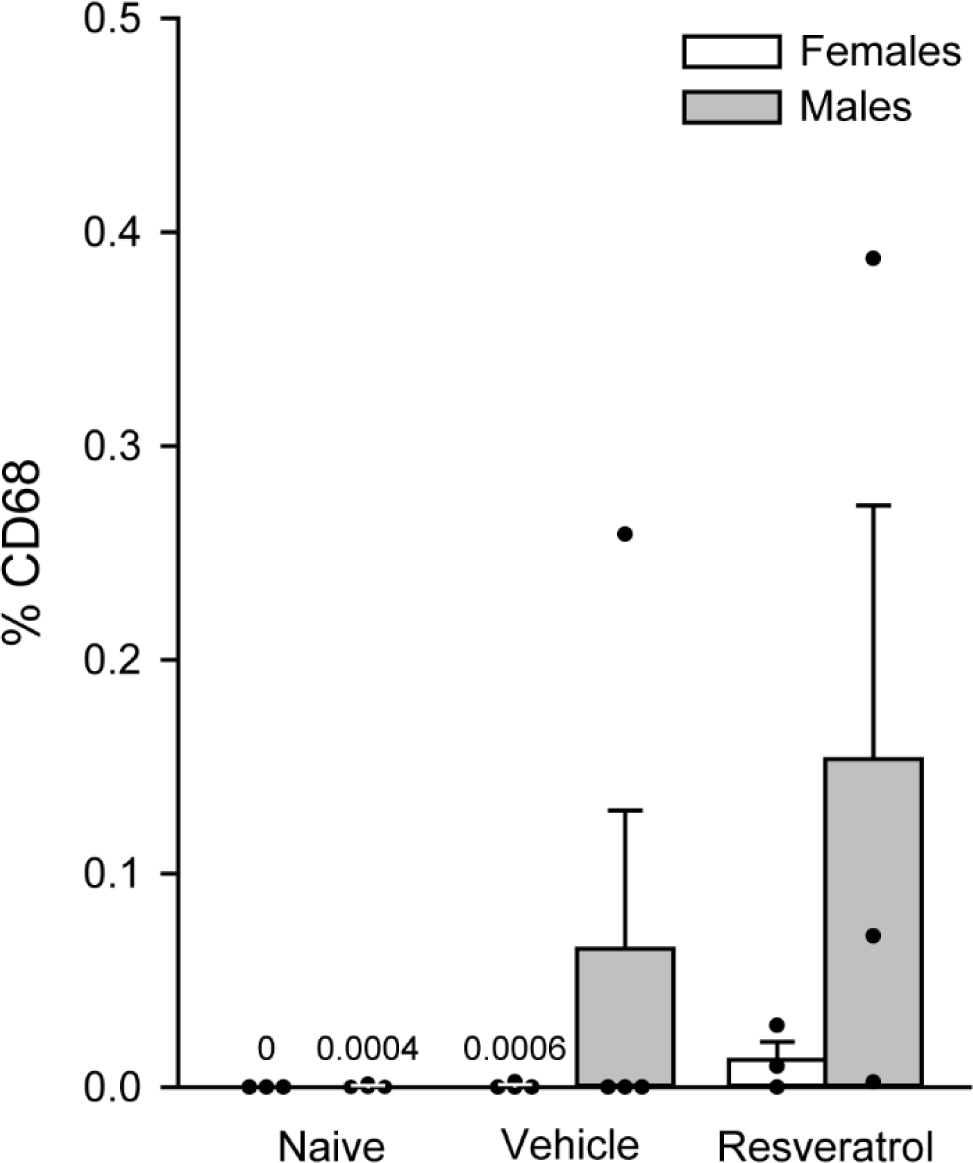
Resveratrol increases CD68 in the corneas of abraded rats. The % CD68 in resveratrol-treated abraded corneas was significantly higher than the % CD68 measured in naïve corneas. Vehicle-treated corneas represented a middle ground between naïve corneas and resveratrol corneas, although there was a trend towards a significant difference between vehicle and resveratrol treatment groups. Non-transformed % CD68 data from individual rats for each group are represented by dots and bars represent mean % CD68 ± SEM. The mean % CD68 for the Naïve group is listed on the graph. % CD68 values for males and females were combined for each treatment group.

## Discussion

This study demonstrates that topical resveratrol caused a sex-specific and dose-dependent increase in tear production but was not an effective analgesic for post-PRK ocular pain in either sex. In fact, resveratrol appeared to have a sex-specific detrimental effect on males by slowing the behavioral recovery after corneal abrasion. Treatment with resveratrol had no effect on epithelial thickness or corneal nerve density, although it increased CGRP expression in corneal nerves in abraded males as compared to abraded females. Resveratrol also increased the density of CD68 in abraded corneal epithelia, although corneal CGRP and CD68 measurements were not directly correlated with the sustained increase in OT behavior in individual abraded males. The underlying mechanism of resveratrol’s effects on tear production and spontaneous pain in injured male rats is unclear. In the current study, we abraded, treated and assessed the left cornea only. It is possible that there were contralateral effects of corneal abrasion (Hegarty et al., 2018) and ocular resveratrol treatment.

Resveratrol is well suited for local or topical applications due to it being highly lipophilic and lacking good systemic bioavailability (Burton et al., 2017). When deciding on the best formulation for applying resveratrol to the ocular surface, several factors were taken into account: we considered different vehicles available for ophthalmic delivery, whether the vehicle itself contained any chemicals that could harm the corneal epithelium or the corneal nerve population, the state of the drug within the vehicle as well as dosing, therapeutic efficiency and time considerations when applying a drug to the ocular surface of the rat. The ophthalmic ointment used in this study was provided in a semi-solid state that melted when it contacted the ocular surface and was evenly distributed over the abraded area of the cornea within the confines of the retaining ring affixed to the cornea. Administration of ophthalmic ointment using anesthetized animals prevented blinking, thereby allowing for longer contact time between the drug and the cornea. Resveratrol was in crystal form within the ophthalmic ointment and demonstrated a dose-dependent effect on tear production and a sex-specific effect on OT behavior after corneal abrasion, supporting the hypothesis that as least some effective concentration was able to reach epithelial tissues and causes sustained effects one week later. There was no effect of resveratrol on epithelial thickness or corneal nerve density as compared to abraded animals that received vehicle. These data demonstrate that this crystalline formulation and dosing protocol provided efficient drug delivery without damage to the corneal nerve population. This ointment formulation avoided the use of solvents to dissolve the drug in a liquid formulation. For example, DMSO is a common solvent used to dissolve drugs, but it has been found to be toxic to ocular structures (Galvao et al., 2014), including the corneal surface. (Bignami et al., 2014).

Clinical studies on post-PRK pain and other vision correction surgeries have documented an increase in acute pain and photophobia in patients. These are self-reporting and based on ordinal scales that rely on the patients to communicate their pain level and photophobia. Pre-clinical rodent studies must rely on animal behavior to convey pain state and photophobia. Our previous studies used the stimulus-evoked eye wipe test (Hegarty et al., 2018;Hegarty et al., 2017), but the ophthalmic ointment formulation appeared to interfere with accurate assessments of acute pain responses (data not shown). In an effort to better model the clinical experience, and to assess spontaneous pain in animals receiving the ophthalmic ointment formulation, we used orbital tightening (OT), one of the metrics of the Rat Grimace Scale (Sotocinal et al., 2011). While there are other components of this scale, including nose and cheek flattening, ear changes and whisker changes, we decided to focus on OT as it was the most relevant to our corneal abrasion model and the most straightforward metric to assess (Harris et al., 2017). We performed our OT assay in ambient light conditions in our behavior room, so it is possible that our spontaneous pain measurements also reflect photophobia (Harris et al., 2017) in the abraded rats, but further studies are needed to confirm. An important advantage of the OT assessment is that it allows for measuring spontaneous pain at several time points, including before and after corneal abrasion. This is in contrast to the stimulus-evoked eye wipe test, which we only measure once in each animal to avoid sensitization and the effects of noxious stimuli on corneal nerve density (Hegarty et al., 2017). Importantly, we found that OT scores of vehicle-treated abraded rats were not different from untreated abraded rats at any time point, demonstrating that ophthalmic ointment had no effect on OT scores that could interfere with our analysis and that any differences we observed were due to the drug.

In our previous study, tear production was significantly decreased 1 week after corneal abrasion (Hegarty et al., 2018). We hypothesized that resveratrol treatment would prevent this decrease in tear production. Unexpectedly, we found that abraded male and female rats that received vehicle ointment demonstrated no change in tear production at 1 week post-abrasion, suggesting that the vehicle itself prevented the abrasion-induced decrease in tear production.

The ophthalmic ointment used as the vehicle in this study is hypertonic and used clinically to reduce corneal inflammation by drawing fluid out of the cornea, (Chow and Chan, 2021) thus it is possible that the ointment drew fluid out of the cornea which prevented or masked a decrease in tear production after abrasion injury. Low concentration resveratrol (1%) also had no effect on tear production. One previous study in female mice found that a low concentration (0.01%) resveratrol liquid eye drop formulation did not prevent a decrease in tear production in a mouse model of dry eye, although it decreased clinical signs of superficial keratitis and inflammation (Abengozar-Vela et al., 2019). Therefore, it is possible that 1% resveratrol was too low to affect a change beyond the vehicle effect. In contrast, higher concentrations of resveratrol significantly increased tear production in males one week after abrasion when compared to baseline measurements, while females gained no benefit from resveratrol treatment. This suggests that females are less susceptible than males to resveratrol’s effects on homeostatic tear production. To our knowledge, this is also the first preclinical study that has demonstrated a dose-dependent sex-specific effect of resveratrol on tear production, although the mechanisms underlying this sex difference are unclear.

In the current study, we used established immunohistochemistry, confocal microscopy and image analysis methods (Hegarty et al., 2018;Hegarty et al., 2017) to measure changes in epithelial thickness, the density of subbasal and intraepithelial corneal nerves, including CGRP-labeled nerves and the infiltration of macrophages. We found that 1 week post-abrasion, epithelial thickness was still significantly reduced in abraded corneas of both sexes as compared to naïve controls, with or without resveratrol. This differs from our previous corneal abrasion study, in which epithelial thickness in untreated abraded corneas was back to naïve levels 1 week after corneal abrasion (Hegarty et al., 2018). There were no differences between vehicle and resveratrol corneas, suggesting that the ophthalmic ointment used as vehicle prevented the full recovery of epithelial thickness after abrasion. As described above, the hypertonic ointment in the current study is used clinically to reduce corneal edema. Many clinical studies have examined the use of hypertonic saline aqueous and ointment formulations to reduce corneal edema in a variety of corneal conditions such as bullous keratopathy, Fuchs endothelial dystrophy, recurrent corneal erosions and post-cataract or keratoplasty surgery (Chow and Chan, 2021). In many cases, topical hypertonic drops and ointment were successful in reducing corneal edema, pachymetrically measured as a decrease in corneal thickness (Chow and Chan, 2021;Yin G.H.W and Levy, 2018). Therefore, it is possible that hypertonic ophthalmic ointment removed fluid from the abraded eye, thereby decreasing corneal epithelial thickness in all treated corneas. Resveratrol treatment had no effect on corneal epithelial thickness. A previous study found that once daily topical application of a resveratrol-loaded gel for 14 days following a corneal alkaline burn promoted recovery of the corneal epithelium in rats (Li M. et al., 2021). The current study only used 3 applications of resveratrol prior to and just after corneal abrasion and looked at one time point after abrasion. Therefore, it is possible that we did not see full epithelial recovery after resveratrol because of the time course and dosing schedule of our study.

Corneal nerve density was reduced in corneas from abraded rats of both sexes as compared to naïve rats 1 week post-abrasion. This decrease at 1 week is consistent with our previous study that found a significant reduction in β tubulin-labeled corneal nerves 1 week after abrasion (Hegarty et al., 2018). Topical resveratrol did not promote recovery of corneal nerve density after abrasion in our study, although some previous studies have implicated resveratrol as a neuroprotective agent based on its actions in restoring sensation after injury (Burton et al., 2017;Hao et al., 2020;Li X. et al., 2019;Tillu et al., 2012;Wang Y. et al., 2020;Yang et al., 2016;Yin Y. et al., 2019). It is possible that the route of administration, dosage, or dosing schedule in our study may not have been sufficient to increase corneal nerve density after injury.

CGRP is a nociceptive marker that is the most abundant peptide in corneal afferents terminating in the trigeminal dorsal horn (Hegarty et al., 2010) and in the trigeminal ganglion cell bodies (LaVail et al., 1993;Nakamura et al., 2007). Our previous study found that the volume of CGRP labeling within β-tubulin-labeled corneal nerves was increased in abraded male rats at 1 week, while the number of CGRP-labeled trigeminal ganglion neurons was similar to unabraded rats (Hegarty et al., 2018). In the current study, we normalized the volume of CGRP in corneal nerves to the volume of the epithelium (Hegarty et al., 2017), instead of the volume of β-tubulin (Hegarty et al., 2018), to account for significant differences in the epithelial volume among the abraded animals as described above. Therefore, a direct comparison of the current % CGRP results to our previous study is not possible. Abraded females had less CGRP in the epithelium (% CGRP), regardless of treatment, while males demonstrated no change in CGRP at one week after abrasion, suggesting that resveratrol has no effect on CGRP after injury. There was a difference in % CGRP between abraded resveratrol males and abraded resveratrol females. It is possible that resveratrol exerted a small effect on % CGRP in males that was perhaps masked by the variance in the data when compared to naïve and abraded vehicle males. We examined whether % CGRP values correlated with OT score, as there was variance in both measures, especially in males. Our analysis demonstrates that female OT scores were tightly positively correlated with % CGRP but male OT scores were not. A sex-specific relationship between peripheral CGRP and pain was demonstrated in a recent study in which dural injections of CGRP decreased facial withdrawal thresholds and increased facial grimace scores in female rats and mice, but not male rodents (Avona et al., 2019). The current study is consistent with a close relationship between CGRP and pain in females, but does not explain the situation in males.

Injury to the corneal epithelium causes an inflammatory response in which macrophages migrate into the wounded area and are essential to the corneal wound-healing process (Bellner et al., 2015;Ljubimov and Saghizadeh, 2015). Previous studies in other systems have demonstrated that resveratrol has anti-inflammatory activity in macrophages and modulates other aspects of the immune response to tissue injury (Malaguarnera, 2019). We used CD68, a pan-macrophage marker, to examine whether resveratrol affected macrophage labeling in the abraded cornea. Resveratrol significantly increased CD68-labeled macrophages in the corneal epithelium of abraded animals compared to naïve corneas, but there was no significant difference compared to vehicle-treated abraded corneas, although there was a trend. When looking at the data for each individual cornea, it is clear that the effects of abrasion and resveratrol treatment were not consistent and demonstrated a great deal of variability in the male abraded corneas, while the % CD68 measured in female corneas were much less variable. The % CD68 from each cornea did not correlate with OT scores, suggesting that the presence of macrophages on the corneal epithelium was not the source of sustained spontaneous pain in males after abrasion. We examined corneal metrics at one time point after abrasion and resveratrol treatment, but it is possible that macrophage infiltration and resveratrol’s effects on macrophages are more pronounced at earlier time points. We did not differentiate between M1- and M2-type macrophages, which are thought to be involved in the initiation and inhibition of inflammatory processes, respectively (Liu J. and Li, 2021). A previous study using a myocardial infarction mouse model found that daily oral resveratrol promoted the proliferation of macrophages in the area surrounding that cardiac infarct and their polarization to M2-type macrophages (Liu S. et al., 2019). It is possible that corneal abrasion and resveratrol treatment induces complex changes in macrophage phenotype that are modulated by time and sex. The current study is consistent with an increase in macrophage expression in injured tissue, but we do not know if macrophages are mediating inflammation or healing.

The mechanisms underlying the sex differences in post-abrasion OT behavior, tear production and corneal % CGRP were not examined in the current study but the influence of gonadal hormones and resveratrol’s weak estrogenic activity (Fremont, 2000;Thaung Zaw et al., 2020)cannot be ruled out. Resveratrol has affinity for estrogen receptors although to a significantly lesser degree than natural estrogen (Qasem, 2020). Resveratrol is thought to be an agonist of the estrogen receptor, but it may also act as an antagonist depending on the dose and the cell system under study (Qasem, 2020;Salehi et al., 2018). Although estrogen is the primary gonadal hormone in females, estrogen and its receptors are also present in lower levels in males (Suzuki et al., 2001;Tachibana et al., 2000;Wickham et al., 2000). Several clinical pain conditions are more prevalent or have a higher degree of severity in females, such as migraine, neuropathic pain, fibromyalgia, osteoarthritis and temporomandibular pain, and there is considerable evidence that these pain conditions are influenced by endogenous and supplemental estrogen (Fillingim et al., 2009;Manson, 2010). Studies in male rodents often involve either systemic or local injection of estrogen or estrogenic supplements similar to resveratrol, and the results range from estrogen being analgesic (Lee J. Y. et al., 2018) to pro-nociceptive (Ji et al., 2018;Qu et al., 2015;Soriano et al., 2019), to having no effect at all (Aloisi et al., 2010). The effects of hormone supplementation on tear production have also been mixed in preclinical studies (Kumar et al., 2018;Song et al., 2014) using ovariectomized female animals to model post-menopausal women with low estrogen levels, similar to males. The presence of estrogen receptors in male corneas (Suzuki et al., 2001;Tachibana et al., 2000;Wickham et al., 2000) and the colocalization of estrogen receptors and CGRP in the trigeminal ganglia neurons of male and female rats (Warfvinge et al., 2020) provide anatomical evidence of a relationship between estrogenic activity and CGRP pain amplification. However, further studies are needed to confirm a functional relationship and a basis for the sex differences observed after topical ocular resveratrol treatment.

Although PRK has been performed for decades, postoperative pain and other ocular symptoms continue to be a challenge for clinicians. Clinical studies of PRK have documented similar increases in postoperative pain and photophobia as measured subjectively by patients on ordinal scales, such as the visual analog scale (VAS) for pain (Colin and Paquette, 2006;Gaeckle, 2021;Palochak et al., 2020;Ripa et al., 2020;Shetty et al., 2019;Zarei-Ghanavati et al., 2019). Clinical signs of dysfunctional tear syndrome or dry eye, as measured with the Schirmer’s test and described by patients as burning, dryness, irritation and the sensation of a foreign body in the eye are also increased in PRK patients (Beheshtnejad et al., 2015;Bower et al., 2015;Quinto et al., 2008). Several clinical studies have been dedicated to finding better drugs, routes of drug administration and effective dosing regimens for PRK patients. For example, topical corticosteroids, oral (ibuprofen, naproxen sodium) and topical (ketorolac, diclofenac) nonsteroidal anti-inflammatory drugs (NSAIDs), narcotics (codeine, oxycodone), and other supplements (amino acids) have been tested with varying degrees of success in PRK patients. The effectiveness of these drugs as analgesics post-PRK needs to be weighed against the side effects that often accompany them. For example, corticosteroids can cause an increase in intraocular pressure and increased risk of developing glaucoma (Javadi et al., 2008;O’Brart et al., 1994), oral NSAIDs can cause gastrointestinal irritation and ulcerations, topical NSAIDs can lead to delayed epithelial wound healing, punctate keratitis and corneal melt (Ripa et al., 2020;Shetty et al., 2019) and narcotics carry the risk of dependence and misuse as well as respiratory depression and gastrointestinal effects (Palochak et al., 2020;Volkow et al., 2019). There is a need for better therapeutics that will address the post-PRK symptom profile and their underlying mechanisms.

Previous *in vivo* rat studies involving abrasion or debridement of the corneal epithelium (Daull et al., 2016;Green et al., 2015;Ho et al., 2013;Kim et al., 2010;Kim et al., 2013;McAlvin et al., 2015;Moore et al., 2013;Nagai et al., 2018;Nagai et al., 2010;Romano et al., 2014;Wang F. et al., 2020;Wang L. et al., 2013;Woodruff et al., 2019), or excimer laser ablation of the anterior stroma (Azar et al., 1996;Hafezi et al., 2018;Lee K. S. et al., 2012;Lu et al., 1999;O’Brien et al., 1998;Prada and Ngo-Tu, 2011;Sandvig et al., 1997;Varela et al., 2002;Wang T. et al., 2013;Woo et al., 2014) have focused on mechanisms of corneal epithelial cell wound healing, and the effects of topical therapeutics after debridement or ablation. These studies were primarily performed in male rats and used fluorescein ophthalmic staining to examine corneal wound healing (Ho et al., 2013;Kim et al., 2010;Kim et al., 2013;McAlvin et al., 2015;Moore et al., 2013;Nagai et al., 2018;Nagai et al., 2010;Romano et al., 2014;Wang F. et al., 2020;Wang L. et al., 2013;Woodruff et al., 2019). Few studies have looked at the impact of topical therapeutics on corneal nerve reinnervation (Romano et al., 2014) and spontaneous ocular behavior (Green et al., 2015) in rats. The similarities between common clinical post-PRK symptoms and the results of our previous study, combined with the results of the current study suggest that our corneal abrasion model and assessments are an accessible and comprehensive strategy to test new therapeutics for post-PRK symptoms. The current study also highlights the importance of using males and females to better understand sex differences in pain processing and analgesic effectiveness.

## Abbreviations

CGRP: calcitonin gene-related peptide

NSAIDs: non-steroidal anti-inflammatory drugs

OT: orbital tightening

PB: phosphate buffer

PRK: photorefractive keratectomy

Res: resveratrol

ROI: region of interest

TS: Tris-buffered saline

Veh: vehicle

## Acknowledgements

The authors thank the OHSU Medicinal Chemistry Core and the OHSU Advanced Light Microscopy Core for their guidance and effort in this study.

## Author Contributions

DMH: performed animal work, immunohistochemistry, confocal imaging and image analysis, writing original manuscript; JRC and DN: analysis of OT behavior; VSH: prepared ophthalmic resveratrol formulation; DIR: advisory role, funding acquisition; TJP and GD: design of study, funding acquisition, advisory role; SAA: design of study, funding acquisition, supervision; data interpretation. All authors were involved with editing the final manuscript.

